# Multivariate Heteroscedasticity Models for Functional Brain Connectivity

**DOI:** 10.1101/154468

**Authors:** Christof Seiler, Susan Holmes

## Abstract

Functional brain connectivity is the co-occurrence of brain activity in different areas during resting and while doing tasks. The data of interest are multivariate timeseries measured simultaneously across brain parcels using resting-state fMRI (rfMRI). We analyze functional connectivity using two heteroscedasticity models. Our first model is low-dimensional and scales linearly in the number of brain parcels. Our second model scales quadratically. We apply both models to data from the Human Connectome Project (HCP) comparing connectivity between short and conventional sleepers. We find stronger functional connectivity in short than conventional sleepers in brain areas consistent with previous findings. This might be due to subjects falling asleep in the scanner. Consequently, we recommend the inclusion of average sleep duration as a covariate to remove unwanted variation in rfMRI studies. A power analysis using the HCP data shows that a sample size of 40 detects 50% of the connectivity at a false discovery rate of 20%. We provide implementations using R and the probabilistic programming language Stan.

## 1 INTRODUCTION

Functional connectivity focuses on the exploration of neurophysiological measures of brain activity between brain regions (Friston, 2011; Smith, 2012; Varoquaux and Craddock, 2013). Functional connectivity studies have increased our understanding of the basic structure of the brain (Sporns et al., 2004; Eguiluz et al., 2004; Bassett and Bullmore, 2006; Fox and Raichle, 2007; Bullmore and Sporns, 2009; Van Den Heuvel and Pol, 2010) and provided insights into pathologies (Greicius et al., 2003; Greicius, 2008; Biswal et al., 2010; Fox and Greicius, 2010).

From the statistical point of view, functional connectivity is the problem of estimating covariance matrices, precision matrices, or correlation matrices from timeseries data. These matrices encode the level of connectivity between any two brain regions. The timeseries are derived from resting-state fMRI (rfMRI) by averaging individual voxels over parcels in the gray matter. We define parcels manually or with data-driven brain parcellation algorithms. The final goal can be an exploratory or a differential analysis comparing connectivity across regions between experimental conditions and time (Preti et al., 2016). Many statistical methods are available to estimate covariance matrices, precision matrices, or correlation matrices from multivariate data. The sample covariance and its inverse, or the the sample correlation matrix are usually poor estimators because of the high-dimensionality of the data (large number of parcels p and small number of subjects). The number of parameters grows quadratically in the number of regions with *p*(*p* − 1)/2 possible pairwise connections between parcels. Therefore more elaborate estimators need to be employed, such as the Graphical Lasso (Friedman et al., 2008) estimator for inverse-covariance matrices or the Ledoit-Wolf shrinkage estimator (Ledoit and Wolf, 2004) for correlation matrices. Application of these methods to rfMRI are available (Varoquaux et al., 2010a,b; Smith et al., 2011; Ryali et al., 2012; Varoquaux et al., 2012).

The estimation of connectivity is usually only the first step and leads to downstream differential analyses comparing connectivity between experimental conditions or between subgroups. For instance, we will compare the connectivity of short sleepers with conventional sleepers available as preprocessed timeseries from the Human Connectome Project (Van Essen et al., 2013). One approach is massive univariate testing of each of the *p*(*p* − 1)/2 connections by linear modeling. Such an approach allows us to test different contrasts and include batch factors or random effect terms (Lewis et al., 2009; Grillon et al., 2013). It lacks statistical power because it ignores possible dependencies between elements in the connectivity matrix. An alternative is to assess selected functionals or summary statistics rather than individual elements in the connectivity matrix (Stam, 2004; Salvador et al., 2005; Achard et al., 2006; Marrelec et al., 2008; Bullmore and Sporns, 2009; Ginestet et al., to appear). Another approach is to flip response variable and explanatory variable and predict experimental condition (or subgroup) from connectivity matrices (or functionals of matrices) through machine learning (Craddock et al., 2012; Dosenbach et al., 2010). These approaches lack interpretability in terms of brain function.

From a statistical viewpoint, the problem boils down to modeling heteroscedasticity. Heteroscedasticity is said to occur when the variance of the unobservable error, conditional on explanatory variables, is not constant. For example, consider the regression problem predicting expenditure on meals from income. People with higher income will have greater variability in their choices of food consumption. A poorer person will have less choice, constrained to inexpensive foods. In functional connectivity, heteroscedasticity is multivariate and variances become covariance matrices. In other words, heteroscedasticity co-occurs among brain parcels and can be explained as a function of explanatory variables.

In this article, we propose a low-dimensional multivariate heteroscedasticity model for functional connectivity. Our model is of intermediary complexity, between modeling all *p*(*p* − 1)/2 connections and only using global summary statistics. Our model builds on the covariance regression model introduced by Hoff and Niu (2012). It includes a random effects term that describes heteroscedasticity in the multivariate response variable. We adapt it for functional connectivity and implement it using the statistical programming language Stan. Additionally, we perform preliminary thinning of the observed multivariate timeseries from *N* to the effective sample size *n*. Using *n* reduces false positives and speeds up computations by a factor of *N*/*n*. To find the appropriate *n*, we compute the autocorrelation as it is common in the Markov chain Monte Carlo literature. We compare our low-dimensional model to a full covariance model contained in the class of linear covariance models introduced by Anderson (1973). Both models are used to analyze real data from HCP comparing connectivity between short and conventional sleepers.

## 2 MATERIAL & METHODS

### 2.1 Data

We analyzed data from the WU-Minn HCP 1200 Subjects Data Release. We focus on the functional-resting fMRI (rfMRI) data of 820 subjects. The images were acquired in four runs of approximately 15 minutes each. Acquisition ranged over 13 periods (Q01, Q02, …, Q13). We separated the subjects into two groups: short sleepers (≤ 6 hours) or conventional sleepers (7 to 9 hours) as defined by the National Sleep Foundation (Hirshkowitz et al., 2015). This results in 489 conventional and 241 short sleepers. The HCP 1200 data repository contains images processed at different levels: spatially registered images, functional timeseries, and connectivity matrices. We work with the preprocessed timeseries data. In particular, the rfMRI preprocessing pipeline includes both spatial (Glasser et al., 2013) and temporal preprocessing (Smith et al., 2013). The spatial preprocessing uses tools from FSL (Jenkinson et al., 2012) and FreeSurfer (Fischl et al., 1999) to minimize distortions and align subject-specific brain anatomy to reference atlases using volume-based and surface-based registration methods. After spatial preprocessing, artifacts are removed from each subject individually (Salimi-Khorshidi et al., 2014; Griffanti et al., 2014), then the data are temporally demeaned and variance stabilized (Beckmann and Smith, 2004), and further denoised using a group-PCA (Smith et al., 2014). Components of a spatial group-ICA (Hyv et al., 1999; Beckmann and Smith, 2004) are mapped to each subject defining parcels (Glasser et al., 2013). The ICA-weighted voxelwise rfMRI signal are averaged over each component. Each weighted average represents one row in the multivariate timeseries. Note that parcels obtained in this way are not necessary spatially contiguous, in particular, they can overlap and include multiple spatially separated regions. HCP provides a range of ICA components 15, 25, 50, 100, 200, and 300. We choose 15 (Figure 1) for our analysis to allow for comparison with prior sleep related findings on a partially overlapping dataset (Curtis et al., 2016).

**Figure 1.**
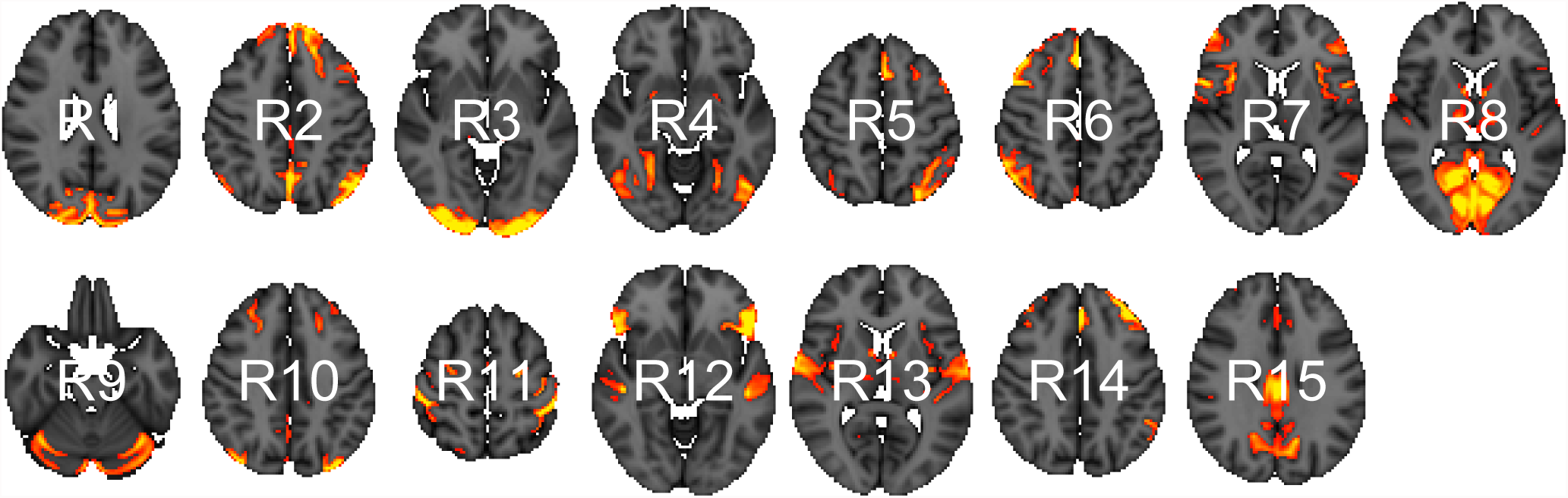
Parcels derived from spatial group-ICA. Created at the most relevant axial slices in MNI152 space. According to Smith et al. (2009), these parcels map to visual areas (R1, R3, R4, and R8), sensorimotor (R7 and R11), cognition-language (R2, R5, R10, and R14), perception-somesthesis-pain (R2, R6, R10, and R14), cerebellum (R9), executive control (R12), auditory (R12 and R13), and default network (R15).

### 2.2 Low-Dimensional Covariance Regression

In this section, we introduce a low-dimensional linear model to compare connectivity between experimental conditions or subgroups.

#### 2.2.1 Model

The data we observe are *p*-dimensional multivariate vectors ***y***_1_,… ***y**_N_*. We assume that the observations are mean-centered so that 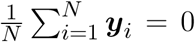. After centering, we subsample each timeseries at *n* < *N* time points to remove temporal dependencies between observations (Section 2.2.2). We are given a set of explanatory variables ***x**_i_* that encode experimental conditions or subgroups, e.g. element one is the intercept 1 and element two is 0 for conventional and 1 for short sleepers. We bind the ***x**_i_*’s row-wise into the usual design matrix ***X***. Our model

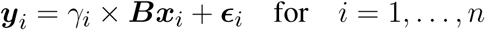

has a random effects term *γ_i_* × ***Bx****_i_* and an independent and identically distributed error term *ϵ_i_*. We suppose the two random variables to have

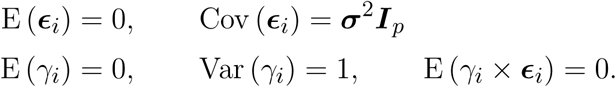

Then, the expected covariance is of the form

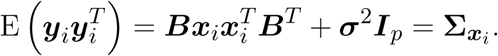

resulting from the inclusion of the random variable *γ_i_*. The covariance matrix **Σ** is indexed by ***x**_i_* to indicate that it changes as a function of the explanatory variables. As with usual univariate linear modeling, we can interpret the coefficients ***B*** as explaining differences between experimental conditions. The matrix ***B*** is *p* × *J* dimensional, where *J* is the number of columns in the *n* × *J* dimensional design matrix ***X***. Here *J* = 2 and the second column encodes the contrast between short sleepers and conventional sleepers. The interpretation of ***B*** is that small values indicate little heteroscedasticity, identical signs indicates positive correlation, and opposite signs indicate negative correlation. For instance, assume that the second column of ***B*** is ***b***_2_ = (−1, 3, 0, 2)*^T^*. The interpretation for these four regions is as follows: region one and two are negatively correlated, so are region one and four, region two and four are positively correlated, and region three is uncorrelated.

The general form of this model was introduced by Hoff and Niu (2012) with the idea of decomposing covariance matrices into covariates explained and unexplained terms. In this original form the unexplained part is parametrized as a full covariance matrix scaling quadratically in the number of regions, i.e. *p*(*p−*1)/*p* parameters. Instead, we parametrize it as a diagonal matrix with independent variance terms for each region. This simplified model scales linearly in the number of regions *p* and can therefore be applied to large brain parcellations.

We use flat priors on both parameters ***σ*** and ***B***. The elements of the ***B*** matrix have a uniform prior on (−∞, ∞), and the elements of ***σ*** vector have a uniform prior on (0, ∞). These priors are improper and do not integrate to one over their support. In case of prior knowledge, it is preferable to use more informative priors. For large *p*, we can add an additional hierarchical level to adjusting for multiple testing by including a common inclusion probability per column in ***B*** (Scott and Berger, 2006; Scott et al., 2010).

As is common in univariate linear modeling, it is possible to encode additional explanatory variables such as subject ID and possible batch factors. It would also be possible to extend the model to include temporal dependencies in the form of spline coefficients. We have not done so here because we wanted to focus explicitly on functional connectivity between regions.

#### 2.2.2 Effective Sample Size

We subsample *n* time points to obtain the Effective Sample Size (ESS). This *n* is smaller than the total number *N* of time points because it accounts for temporal dependency. We propose a procedure to automatically choose *n* using an autocorrelation estimate of the timeseries. This is current practice in the field of Markov chain Monte Carlo and implemented in R package coda (Plummer et al., 2006). The ESS describes how much a dependent sample is worth with respect to an independent sample of the same size. Kass et al. (1998) define ESS via the lag *t* autocorrelation Corr 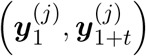 as

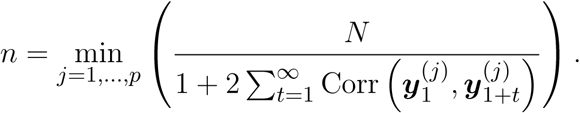

This is a component-wise definition and we follow a conservative approach by taking the minimum over all *p* components as the overall estimator. Intuitively, the larger the autocorrelation the lower is our ESS because we can predict future form current time points. A convenient side-produce of subsampling is reduced computational costs.

#### 2.2.3 Inference

We implement our model in the probabilistic programming language Stan (Carpenter et al., 2016) using R. Stan uses Hamiltonian Monte Carlo to sample efficiently from posterior distributions using automatic differentiation. It removes the need for manually deriving gradients of the posterior distributions, thus making it easy to extend models. Our Stan code is available in our new R package CovRegFC from our GibHub repository. Alternatively, using conjugate priors it is possible to derive a Gibbs sampler to sample from the posterior distribution of a related model as in Hoff and Niu (2012). However, this makes it harder to extend the model.

Due to the non-identifiability of matrix ***B*** up to random sign changes, ***B*** and −***B*** corresponding to the same covariance function, we need to align the posterior samples coming from multiple chains. A general option is to use Procrustes alignment. Procrustes alignment (Korth and Tucker, 1976) is a method for landmark registration (Kendall, 1984; Bookstein, 1986) in the shape statistics literature and an implementation is available in the R package shape (Dryden and Mardia, 1998).

### 2.3 Full Covariance Regression

In this section, we introduce a full covariance linear model.

#### 2.3.1 Model

Here we do not subsample and deal with temporal dependencies in a different way. In this model, the number of observations are the number of subjects *k* = 1, …, *K*. After column-wise centering of each *N* × *p* (recall that *N* is the total number of time points) timeseries ***Y***_1_, …, ***Y****_K_*, we compute sample covariance matrices for each subject 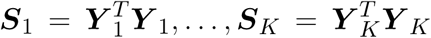. We take this as our “observed” response. Additionally, we have one explanatory vector ***x***_1_, …, ***x**_n_* for each response covariance matrix. In our HCP data subset, we have 730 subjects, so *K* = 730 and we have *K* data point pairs (***S***_1_, ***x***_1_), …, (***S***_*K*_, ***x***_*K*_). We assume that the explanatory vector has two elements: the first element 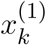 representing the intercept and is equal to one, and the second element 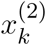 is one for short and zero for conventional sleepers. Our regression model

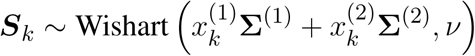

decomposes the “observed” covariance matrix into an intercept term and a term encoding the functional connectivity between sleepers. The second parameter in the Wishart distribution describes the degrees of freedom and has support (*p* − 1, ∞).

We will now describe how to draw samples from the Wishart distribution, this will give us a better intuition for the proposed model. Matrices following a Wishart distribution can be generated by drawing vectors ***y***_1_,…, ***y**_N_* independently from a Normal (0, Σ), storing vectors in a *N* × *p* matrices ***Y***_*i*_, and computing the sample covariance matrix 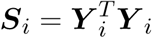. Then, the constructed ***S***_*i*_’s are distributed according to a Wishart distribution with parameters Σ and degrees of freedom *N*. If the ESS is smaller than *N* it will be reflected in the degrees of freedom parameter *ν*. In our model, we will estimate *ν* from the data. In this way, we account for the temporal dependencies in the timeseries. The marginal posterior distribution of *ν* will be highly concentrated around a small degree of freedom (close to *p*) for strongly dependent samples and concentrated around a large degree of freedom (close to *N*) for weakly dependent samples.

To complete our model description, we need to put priors on covariance matrices and the degrees of freedom. We decompose the covariance prior into a standard deviation ***σ*** vector and a correlation matrix Ω for each term

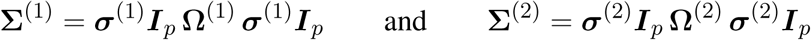

and put a Lewandowski, Kurowicka, and Joe (LKJ) prior on the correlation matrix (Lewandowski et al., 2009) independently for each term

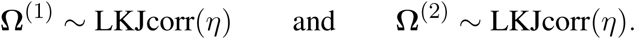

This correlation matrix prior has one parameter *η* that defines the amount of expected correlations. To gain intuition about *η*, we draw samples from the prior for a range of dimensions and parameter settings (Figure 2). The behavior in two dimension is similar to a beta distribution putting mass on either the boundary of the support of the prior or in the center. As we move toward higher dimensions, we can see that the distribution is less sensitive to the parameter *η*. For our model, we set *η* = 1 to enforce a flat prior. We complete our prior description by putting independent flat priors on both the vector of standard deviations ***σ*** and the degrees of freedom *ν*, i.e. uniform prior on (0, ∞) and uniform prior on (*p* − 1, *N* − 1), respectively.

**Figure 2.**
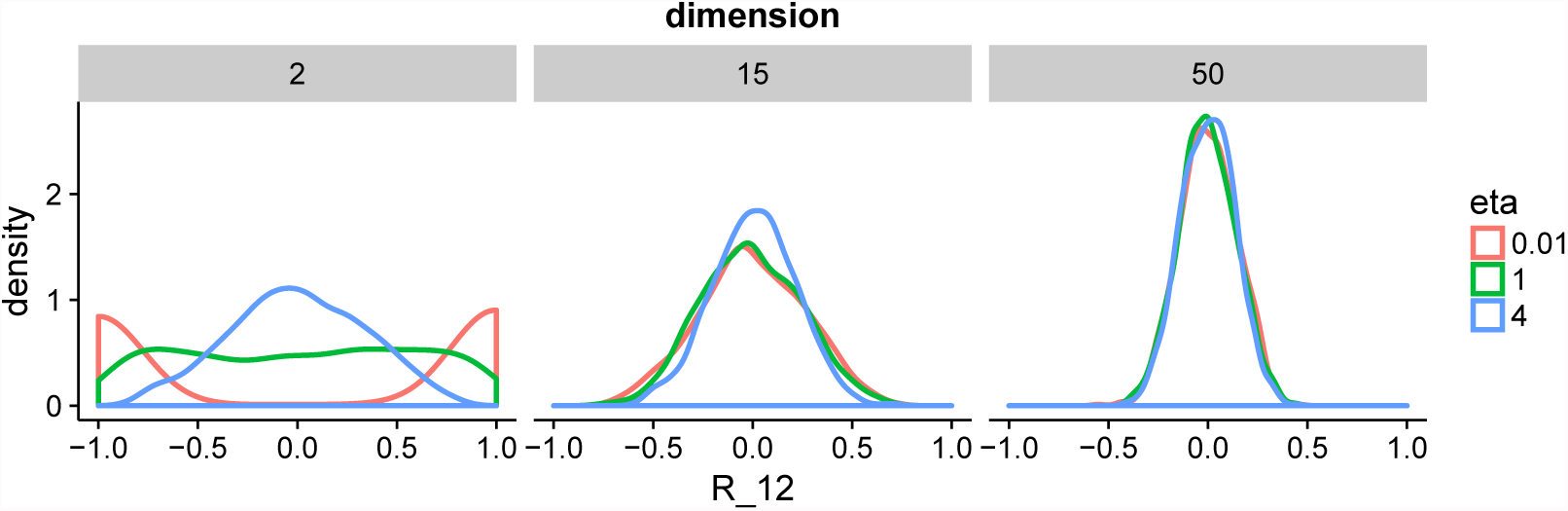
Distribution of 1000 off-diagonal elements *R*_12_ extracted from correlation matrices drawn from the **LKJ** prior. This prior is symmetric, so the distribution will be similar for other off-diagonal elements.

#### 2.3.2 Inference

The number of parameters in the model scales quadratically in the number of regions making this model applicable in the classical statistical setting where we have larger sample sizes than number of predictors. In Section 3.1, we will show an application to the HCP data with *K* = 730 subjects and *p* = 15 regions. Note, Hoff (2009) devised a Gibbs sampler for a similar model using an eigenmodel for the subject-level covariance matrices.

#### 2.3.3 Posterior Analysis and Multiplicity Control

After drawing samples from the posterior, we can evaluate the marginal posterior distributions of standard deviations ***σ***, correlations **Ω**, and degrees of freedom *v*. As mentioned, we assume that the second element in the explanatory vector encodes whether a subject is a short or a conventional sleepers. In this setting, **Ω**^(2)^ represents the difference in correlation between short and conventional sleepers. As we have the marginal posterior distribution for every 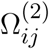, we can evaluate the probability

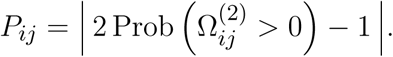

Our interpretation in terms of connectivity is as follows: If *P_ij_* is zero then the correlation is equally probable to be negative or positive. In this case, we are unable to clearly classify the sign of the correlation difference as negative or positive. If *P_ij_* is close to one then the correlation is more probable to be either negative or positive. In this case, we can say that parcel *i* can be seen to be differentially connected to parcel *j*.

There are *p*(*p* − 1)/2 pairwise correlations and we wish to find correlations that are different between the two groups. If the probability *P_ij_* is large, we will report the connection as significantly different. To control for multiple testing, we declare correlations only as significant if they pass a threshold λ. We choose λ to control the posterior expected FDR (Mitra et al., 2016)

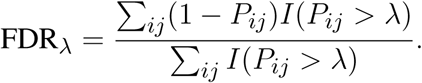

We find λ through grid search for a fixed FDR. This allow us to report only correlations that survive the threshold at a given FDR.

## 3 RESULTS

The HCP released a dataset with 820 timeseries of normal healthy subjects measured during resting-state fMRI (rfMRI). The imaging data is accompanied by demographic and behavioral data including a sleep questionnaire. Approximately 30% Americans are reported short sleepers with 4 to 6 hours of sleep per night. The National Sleep Foundation recommends that adults sleep between 7 to 9 hours. We use both models to analyze the HCP data on 730 participants (after subsetting to short and conventional sleepers) to elucidate difference in functional connectivity between short and conventional sleepers. As mentioned before, the design matrix ***X*** has an intercept 1 and a column encoding short sleepers 1 and conventional sleepers 0, i.e. conventional sleepers are the reference condition. We use a burn-in of 500 steps during which Stan optimizes tuning parameters for the HMC sampler, e.g. the mass matrix and the integration step length. After burn-in, we run HMC for additional 500 steps. To check convergence, we assess traceplots of random parameter subsets. We obtain an effective sample size of 167 for the 15 regions ICA-based parcellation. We now analyze the marginal posterior distribution of each of the parameters.

### 3.1 Differential Analysis

In Figure 3, we summarize and visualize the marginal posterior distribution of the second column in ***B***. In the center part of the plot, we show the posterior distribution as posterior medians (dot) and credible intervals containing 95% of the posterior density (segments). The credible intervals are Bonferroni corrected by fixing the segment endpoints at the 0.05/15 and (1 − 0.05/15) quantiles. Care has to be taken when interpreting the location of segments with respect to the zero coefficient line (red line). Due to the sign non-identifiability of ***B***, we have to ignore on which side the segments are located. Recall that regions on the same side are positively correlated, regions on opposite sides are negatively correlation, and regions overlapping the red line are undecided. To relate the region name back to the anatomy, we plotted the most relevant axial slice in the MNI152 space on the left and the right of the coefficient plot, depending on their sign, respectively. We can make the following observations: Parcels in set 1 (R4, R5, R7, and R9) are positively correlated. Keep in mind that the sign of the coefficient carries no information about the sign of the correlation. So, even though the coefficients are negative the correlations are positive, because they are on the same side of the red line. Parcels in set 2 (R1-R3, R8, R10-R13, and R15) are also positively correlated, for the same reason as before. In contrast, the two parcel sets are negatively correlated, because they are on opposite sides. The connectivity of R6 and R14 are not different from conventional sleepers because their credible intervals overlap the red line. According to the meta analysis in (Smith et al., 2009), parcel set 1 is associate with visual, cognition-language, sensorimotor areas, and the cerebellum; and parcel set 2 with visual, cognition-language, auditory areas, and the default network.

**Figure 3.**
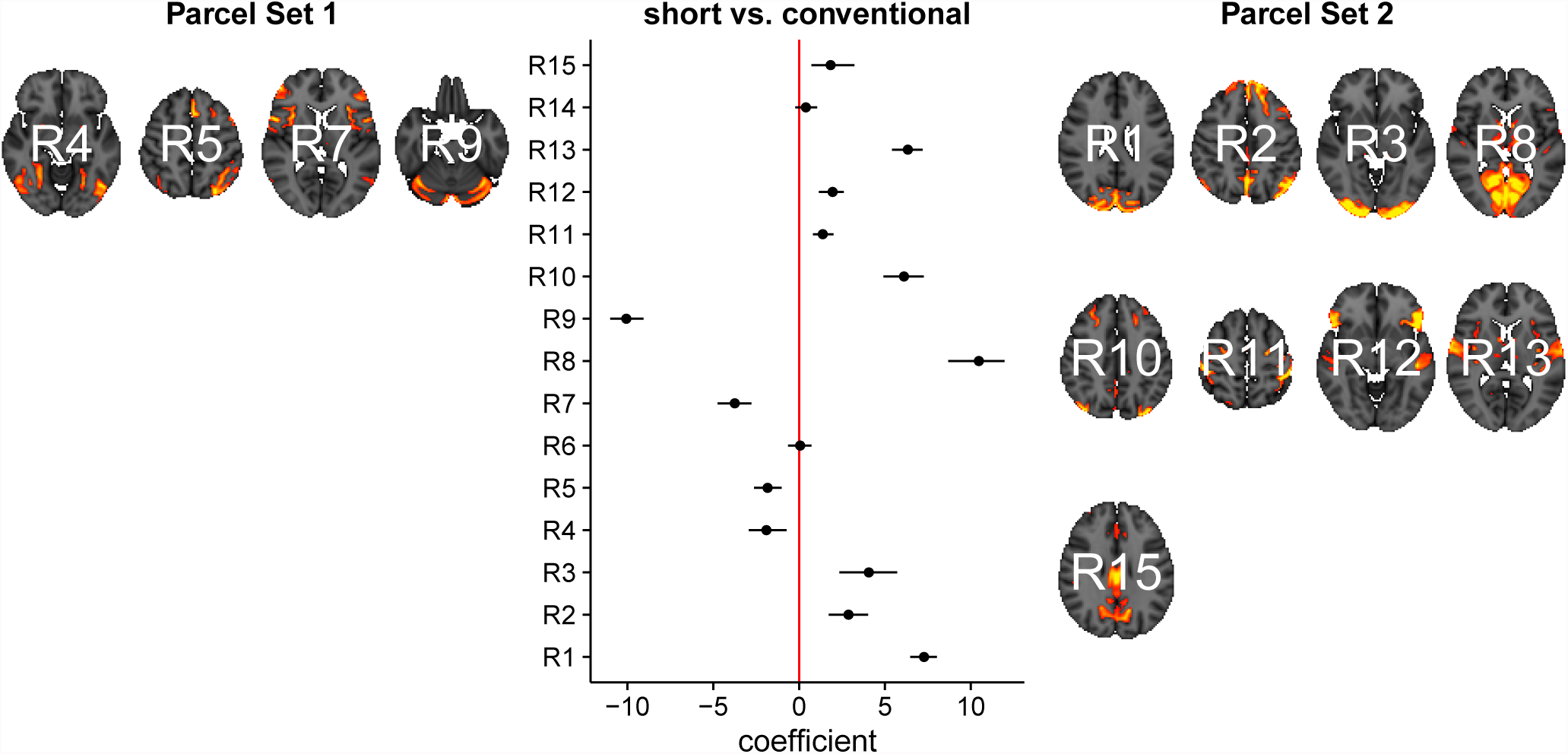
Reduced covariance model. This is the second column in the design matrix encoding the contrast between short and conventional sleepers. The sign is not identifiable; it only matters whether parcels are on the same or opposite side. If they are on the same side, then they are positively correlated. If they are on the opposite side, then they are negatively correlated. The posterior credible intervals are widened according to the number of regions or channels in the plot using the Bonferroni procedure.

We now compare the result from the low-dimensional model with results form the full model. First, we compute the posterior marginal mean of the standard deviations vector *σ*^(2)^ and the correlation matrix magnitude |Ω^(2)^| encoding the difference between short and conventional sleepers (Figures 4). The standard deviation plot on the right shows that parcel R3 varies the most, and that region R2 varies the least. The magnitude correlation plot on the left shows that parcel pair R9 and R13 exhibit the strongest correlation. This is consistent with the low-dimensional model results, where R9 and R13 are in opposite parcel sets. Also these parcels have large effect sizes in the low-dimensional results. In Figure 5, we assess the significance of differential correlations. The color code indicates different FDR levels. Overall strong differences in the correlation structure are visible with a large portion of connections at an FDR of 0.001. In contrast to the low-dimensional model, these are differences in correlations and not whether they are more positively or more negatively correlated.

**Figure 4.**
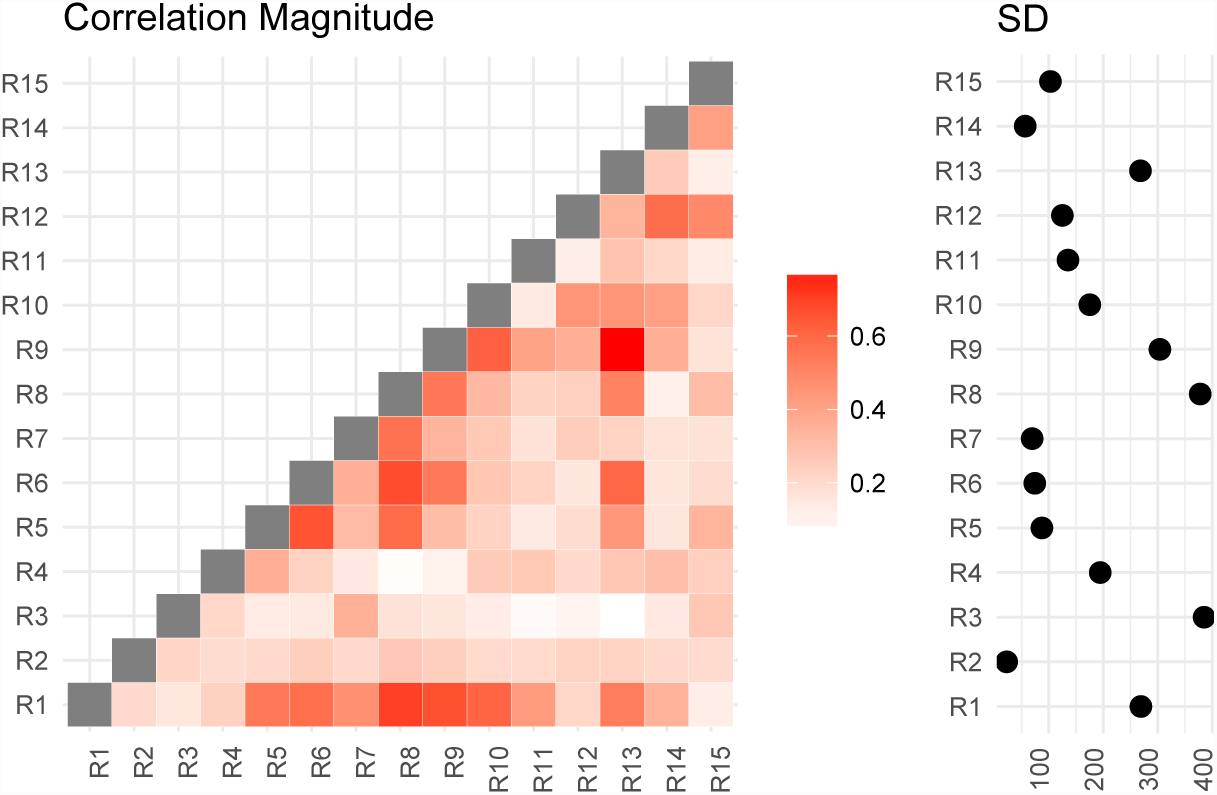
Posterior mean correlations magnitude and standard deviations of the difference between short and conventional sleepers.

**Figure 5.**
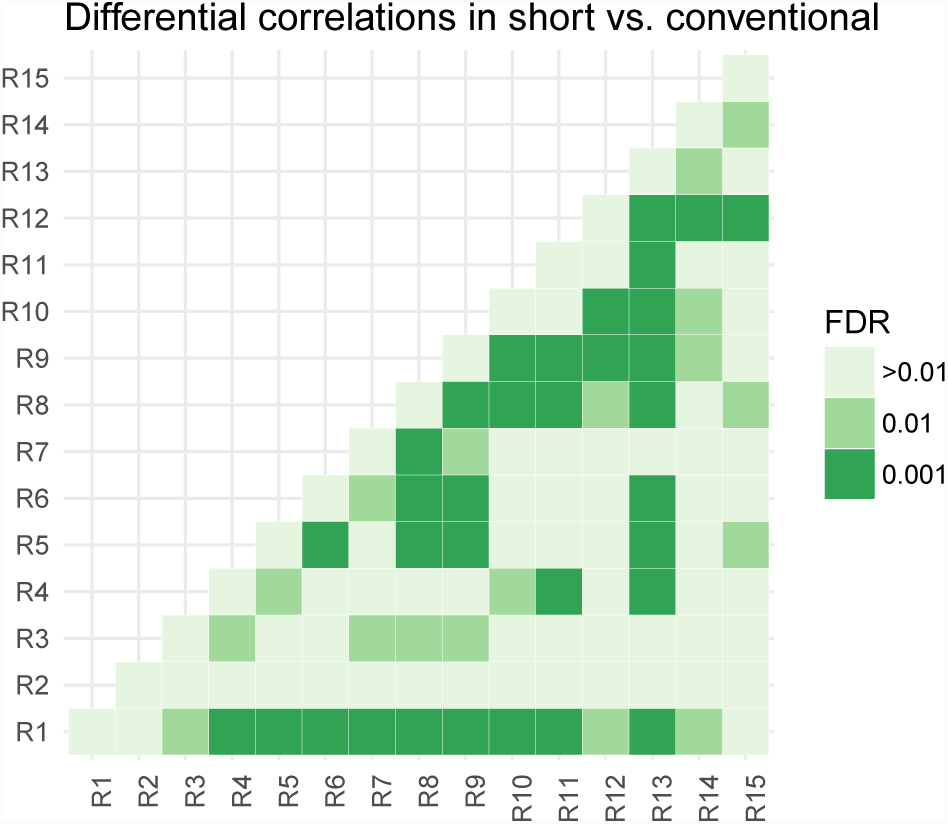
Thresholded connectivity matrix showing the level of differential correlation between all pairs of parcels in short vs. conventional sleepers. Thresholding is chosen to control for posterior expected FDR at three different levels > 0.01, 0.01, and 0.001.

#### 3.1.1 Note on Computation Time

For the low-dimensional model and the available 730 subjects, the computation time for the HMC sampler is around 20 hours on a single core on a modern CPU. For a subsample of 40 subjects, the computation time is around 20 to 25 minutes, and for 80 subjects around 50 to 55 minutes. It is possible to run more chains in parallel to increase the sample size. To combine each run, we need to align the posterior samples using Procrustes alignment as indicated in the methods section.

The full model takes about one hour on a single core, and we run four chains in parallel to increase sample size.

### 3.2 Power Analysis

We design a power analysis (Figure 6) for low-dimensional covariance regression with 15 parcels. As the population we take the available 730 subjects in the HCP data repository that are either short or conventional sleepers and have preprocessed timeseries. We sample 100 times from this population keeping the same ratio between the number of observations for each group, i.e. two thirds conventional and one third short sleepers. We report the average True Positive Rate (TPR) and the False Discovery Rate (FDR) over the 100 samples. The TPR measures the power of our procedure to detect true correlation differences. We count a connection as detected if it is correctly classified as positive or negative correlation. The FDR measures the amount of mistakes we make. The tradeoff between the two can be controlled through the significance level *α*. Power increases linearly with sample size. FDR decrease linearly but at a lower rate with sample size. At samples size 40, we have a power of 50% with an FDR of 20%. This improves to a power of 80% with an FDR of 10% at sample size 160.

**Figure 6.**
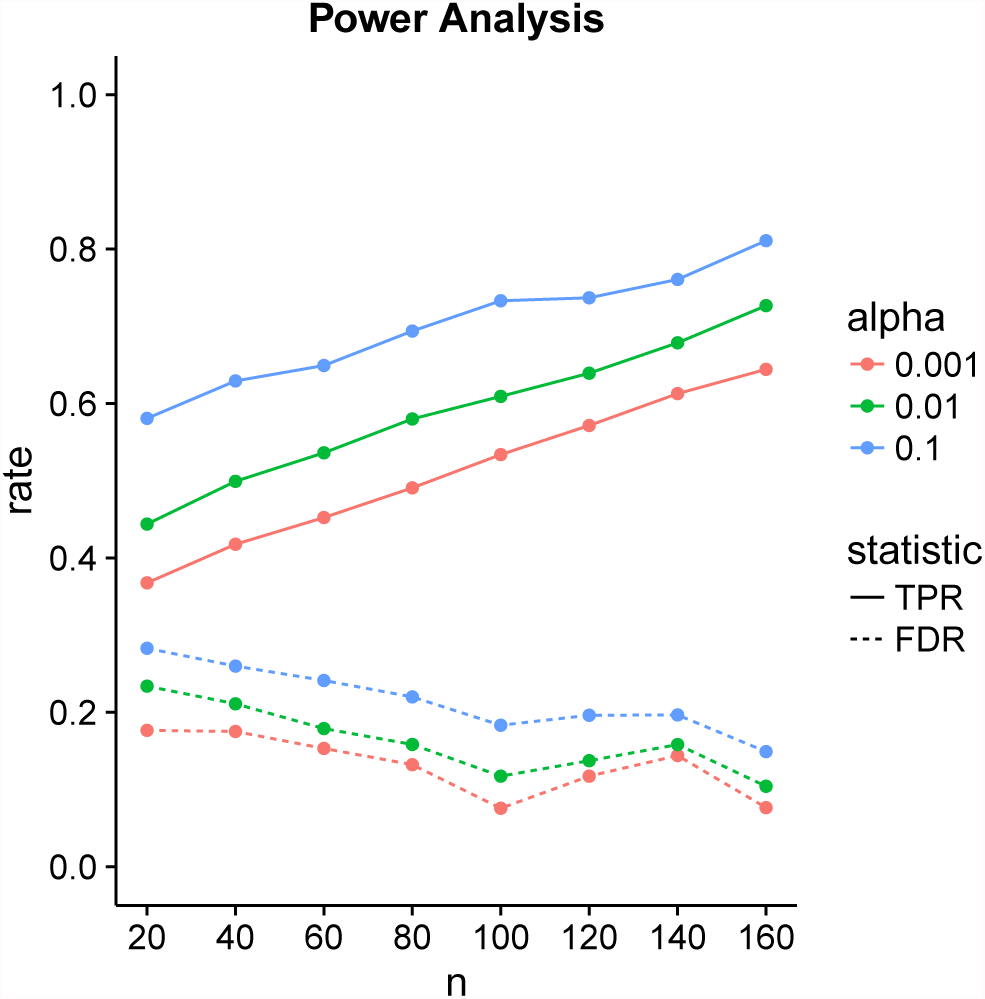
Power analysis for low-dimensional covariance regression with 15 parcels. The two statistics are the True Positive Rate (TPR) and the False Discovery Rate (FDR). The significance level is denoted by α. Points are averages computed over 100 samples from the population.

## 4 DISCUSSION

We introduced two new models for functional connectivity. In particular, the low-dimensional covariance model is able to discover 50% of the correlation differences at a FDR of 20% in a sample size as little as 40. Our Stan implementations make it easy for others to extend our models. We applied both models to the HCP data subset to compare functional connectivity between short and conventional sleepers. Our findings are consistent with Curtis et al. (2016) and Killgore et al. (2012) reporting increases in functional connectivity in short sleepers for primary auditory, primary motor, primary somatosensory, and primary visual cortices. A similar neural signature was observed in experiments examining the transition from resting wakefulness to sleep onset using EEG and rfMRI (Larson-Prior et al., 2009; Tagliazucchi and Laufs, 2014; Davis et al., 2016). Therefore, we recommend the inclusion of the average sleep duration of a subject as a “batch” covariate in the experimental design of rfMRI studies.

A main challenge in covariance regression is the positive definiteness constraint. A solution is to transform the covariance estimation problem into an unconstrained problem thus making estimation and inference easier (Pourahmadi, 2011). One possible transformation starts with a spectral decomposition where the covariance matrix is decomposed into a diagonal matrix of eigenvalues and an orthogonal matrix with normalized eigenvectors as columns. The procedure continues with a global log-transformation to the covariance matrix, which results in a log-transformation of individual eigenvalues and removes the constraint. However mathematically and computationally tempting this approach seems, it remains hard to interpret the log-transformations statistically (Brown et al., 1994; Liechty et al., 2004). An alternative transformation uses a Cholesky decomposition of the covariance matrix. For the Cholesky decomposition, we need a natural ordering of the variables not known a priori for functional connectivity data – a natural ordering could be given if the chronology is known.

Modeling of covariance matrices builds on important geometrical concepts and the medical image analysis community has made significant progress in terms of mathematical descriptions and practical applications motivated by data in diffusion tensor imaging (Pennec, 1999; Moakher, 2005; Pennec, 2006; Arsigny et al., 2006/07; Lenglet et al., 2006; Fletcher and Joshi, 2007; Fillard et al., 2007; Schwartzman et al., 2008; Dryden et al., 2009). The underlying geometry is called Lie group theory and it appears when we consider the covariance matrices as elements in a non-linear space. The matrix log-transformation from the previous paragraph maps covariance matrices to the tangent space where unconstrained operations can be performed; for instance we create a mean by simple elementwise averaging. After computing the mean in tangent space, this mean is mapped back to the constrained space of covariance matrices. Despite the mathematical beauty and algorithmic convenience, statistical interpretations are still unwieldy. However, this does provide a fundamental geometric formulation and enables the use of handy geometrical tools (Absil et al. (2008) for a book-length treatment).

We approach the problem from a statistical viewpoint and frame functional connectivity in terms of modeling heteroscedasticity. This allows us to take advantage of the rich history in statistics and led us to the covariance regression model introduced by Hoff and Niu (2012). We simplify the model to meet the large *p* requirement in neuroscience. The running time for 500 posterior samples on 80 subjects is less than an hour on a single core. This makes our approach applicable to many neuroimaging studies. For larger studies, such as the HCP with 730 subjects, further speed improvements using GPU’s are desirable to reduce computation time.

One possible future application is functional Near-Infrared Spectroscopy (fNIRS), which has gained in popularity due its portability and high temporal resolution. A common approach is to set up a linear model between brain responses at channels locations (Huppert et al., 2009; Ye et al., 2009; Tak and Ye, 2014) and experimental conditions. Thus, our models apply to fNIRS experiments. An additional challenge in fNIRS experiments is channel registration across multiple participants (Liu et al., 2016). Connectivity differences could be due artifacts created by channel misalignments not biology. In the absence of structural MRI, we could add an additional hierarchical level in our low-dimensional model to handle measurement errors accounting for possible misalignments between channels.

We use a conservative component-wise estimate of the ESS. Less conservative multivariate estimators (Vats et al., 2015) might be able to increase statistical power at the cost of an increase in the false discovery rate.

It is possible to append more columns in the design matrix to encode batch factors and subject-specific variability by binding one column per level. In addition to categorical variables, the covariance regression model can handle continuous variables such as head-motion measurement made using an accelerometer. Adding covariates to explain unwanted variation in the data can move some of the preprocessing steps to the functional connectivity analysis step. Such joint modeling can enable the propagation of uncertainty to the downstream analyses.

## REPRODUCIBILITY AND SUPPLEMENTARY MATERIAL

The entire data analysis workflow is available on our GitHub repository:

- https://github.com/ChristofSeiler/CovRegFC_HCP

We also provide a new R package CovRegFC with Stan code:

- https://github.com/ChristofSeiler/CovRegFC

Data preparation and statistical analyses are contained in Rmd files:

- `Low_Dimensional.Rmd`
- `Full.Rmd`
- `Power.Rmd`

By running these files all results and plots can be completely reproduced as html files:

- `Low_Dimensional.html`
- `Full.html`
- `Power.html`

The HCP data is available here:

- https://www.humanconnectome.org/data/

## ACKNOWLEDGMENTS

CS was partially funded by two Swiss NSF postdoctoral fellowships 146281 and 158500. SPH was partially supported by NSF DMS grant 1501767. We thank the NIRS lab at the Center for Interdisciplinary Brain Sciences in the Stanford School of Medicine for introducing us to functional neuroimaging data in the context of fNIRS experiments.

Data were provided by the Human Connectome Project, WU-Minn Consortium (Principal Investigators: David Van Essen and Kamil Ugurbil; 1U54MH091657) funded by the 16 NIH Institutes and Centers that support the NIH Blueprint for Neuroscience Research; and by the McDonnell Center for Systems Neuroscience at Washington University.

## AUTHOR CONTRIBUTIONS

CS wrote an initial draft, performed and implemented the statistical analysis. SPH wrote the final manuscript and provided statistical tools.

